# Assessment of Storm Impact on Coral Reef Structural Complexity

**DOI:** 10.1101/2022.12.04.519015

**Authors:** Matan Yuval, Naama Pearl, Dan Tchernov, Stephane Martinez, Yossi Loya, Avi Bar-Massada, Tali Treibitz

## Abstract

Extreme weather events are increasing in frequency and magnitude. Consequently, it is important to understand their effects and remediation. Resilience reflects the ability of an ecosystem to absorb change, which is important for understanding ecological dynamics and trajectories. To describe the impact of a powerful storm on coral reef structural complexity, we used novel computational tools and detailed 3D reconstructions captured at three time points over three years. Our data-set ***Reefs4D*** of 21 co-registered image-based models enabled us to calculate the differences at seven sites over time and is released with the paper. We employed six geometrical metrics, two of which are new algorithms for calculating fractal dimension of reefs in full 3D. We conducted a multivariate analysis to reveal which sites were affected the most and their relative recovery. We also explored the changes in fractal dimension per size category using our cube-counting algorithm. Three metrics showed a signicant difference between time points, i.e., decline and subsequent recovery in structural complexity. The multivariate analysis and the results per size category showed a similar trend. Coral reef resilience has been the subject of seminal studies in ecology. We add important information to the discussion by focusing on 3D structure through image-based modeling. The full picture shows resilience in structural complexity, suggesting that the reef has not gone through a catastrophic phase shift. Our novel analysis framework is widely transferable and useful for research, monitoring, and management.

**Graphical Abstract:** 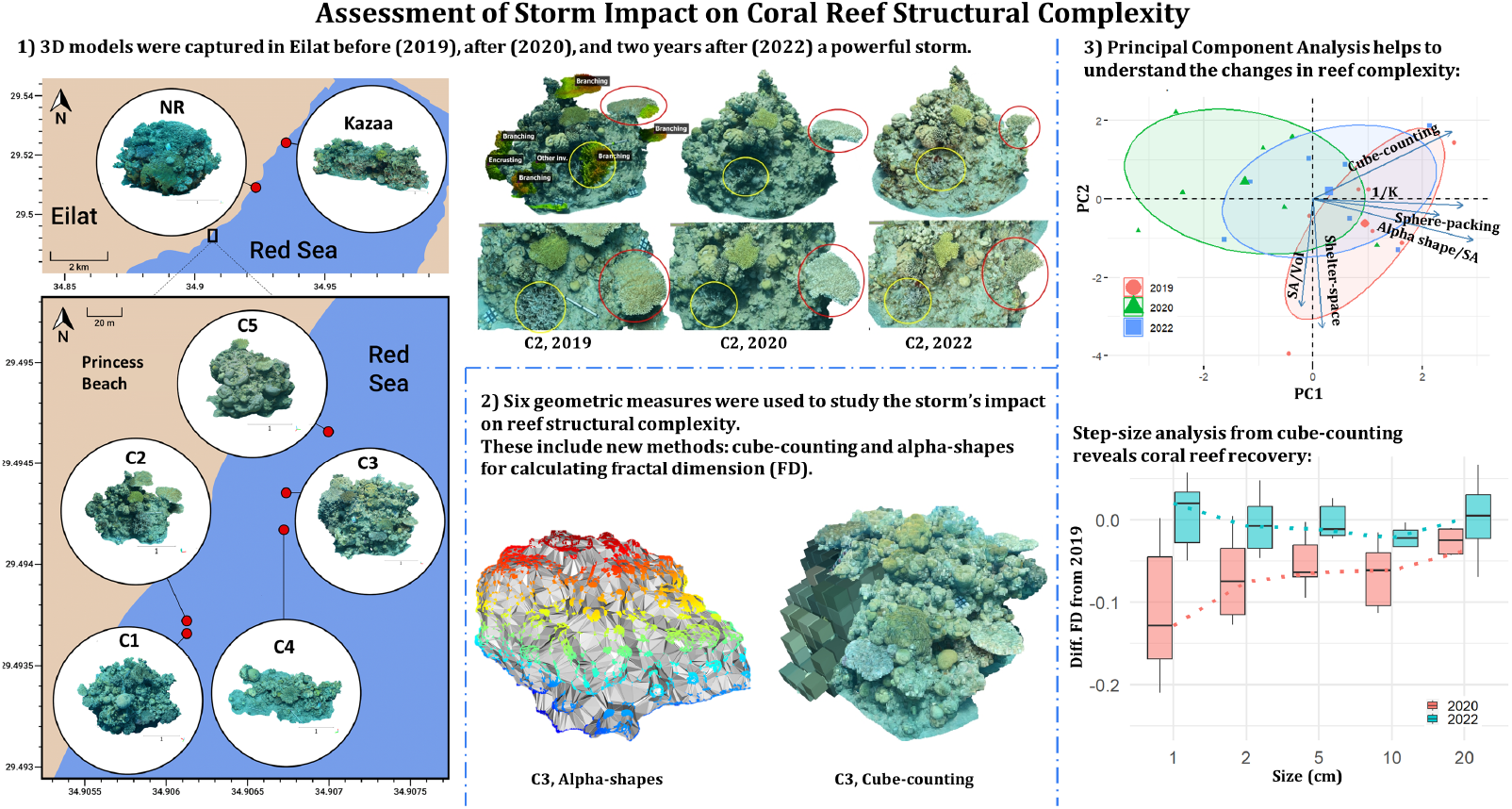

**Highlights:**

- The effect of storms on coral reef 3D structure is poorly understood.
- We studied the impact of a storm on coral reefs using a novel 3D imaging framework.
- We used geometrical metrics including new methods for calculating fractal dimension.
- The reefs recovered in two years with regards to structural complexity.
- Based on 3D analysis the reefs exhibit structural resilience.

## 1 INTRODUCTION

Coral reefs are some of the most important ecosystems on the planet. However, they are undergoing major negative changes due to human activities (Hughes et al. 2003; Hoegh-Guldberg and Bruno 2010; Hughes et al. 2017). Developing and integrating new technologies for coral-reef research and monitoring is key to alleviating the effort of studying these changes in detail across scales, and managing them effectively. Three-dimensional (3D) photogrammetry is increasingly being used to study and monitor coral reefs, but so far it is not exploited to its full potential, i.e., there is still significant underlying information in these models that is not currently being used.

Here we evaluate the impact of a storm on coral reef structural complexity via automated analysis of 3D models. Resilience is defined as the amount of disturbance that an ecosystem can absorb without changing self-organized processes and structures (Holling 1973; Gunderson 2000). This raises the question: what are the stable states (processes and structures) in a transient system, e.g., coral reefs (Standish et al. 2014)? Although many studies have focused on coral reef resilience (Dudgeon et al. 2010; Nyström and Folke 2001), much remains to be learned on the response of reefs to extreme weather events and major storms (Roff and Mumby 2012). Storms can have mixed effects on reefs such as a decrease in live cover of more competitive members, leading to increased diversity (Rogers 1993). Hurricanes have been shown to reduce the cover of dominant fragile forms and to generate space on the reef (Porter et al. 1981). It was even shown that cyclones can contribute to the maintenance of diversity by preventing monopolization and increasing habitat fragmentation (Harmelin-Vivien 1994). However, some of the most negative effects of storms are changes in the topographic structure of the reef, which can lead to an ecosystem collapse through various cascaded pathways.

An ecological system includes a plethora of forces interacting on various spatial, temporal, and organizational levels, forming an intricate web of connections. Resilience is the ability of this web to buffer disturbances. Phase shifts in coral reefs have been most commonly described as an irreversible transition to algae-dominated reefs. However, it has been largely suggested that coral reef environments can capacitate several stable states, representing a more reversible and flexible view (Knowlton 1992; Hughes 1994; Crisp et al. 2022). Although the question of alternate states vs phase shifts is scale and process-dependant, we found that the concept of resilience is useful for our study because its role in both approaches is similar-increased resilience decreases the likelihood of transitions (phase shifts or alternate stable states) (Dudgeon et al. 2010).

The key components of resilience are those that increase the systems’ buffer capacity. Functional richness and diversity enhance the coral reef ecosystem by providing redundancy. Moreover, they enable the emergence of novel ecological configurations which can help to maintain critical processes during or after a disturbance (Paperin et al. 2011). The reef edifice is built by corals and coral diversity increases niche-partitioning and specializations within and across organizational levels. While the coral community may shift following a disturbance, the reef may still be able to provide habitat for fish communities, which in turn perform unique services to the system such as grazing and predation. Again, the question of resilience is focus-dependant as one sub-part of the reef may change while another remains stable, and even the boundaries of fluctuations are often hard to define. Therefore, there is room to examine the resilience of each part separately towards building a comprehensive picture of reef resilience. For example, structural resilience can be viewed as the amount of disturbance that the reef can absorb without losing structural complexity, or more importantly, the reef’s ability to regain structural complexity after a disturbance (Connell and Sousa 1983).

Structural complexity is a major indicator of the reef’s state and an important characteristic of reef resilience. It encapsulates the number of structural features, their sizes, and their distribution. An elaborate and complicated reef framework provides legacy in the case of disturbance (Nyström and Folke 2001). Moreover, in coral reefs structural complexity and biodiversity are clearly linked (Graham and Nash 2013; Ferrari et al. 2016; Alvarez-Filip et al. 2009). While traditional research on coral reef resilience has dealt mainly with community composition metrics, these are difficult to obtain from image-based surveys as they require taxonomic classification. In contrast, structural complexity can be extracted automatically making it more scalable for reef resilience research. Nevertheless, different facets of reef structure can be expressed by different metrics similarly to community metrics, i.e., there is more than one measure for diversity, likewise for structural complexity. In this work, we aim to provide a versatile framework for temporal comparisons of reef 3D structures using several measures. While structural complexity is not interchangeable with resilience, we use it as a proxy for reef-state pointing towards structural resilience.

Fractals are self-similar structures that have gained traction across science and industry since the seminal work of Mandelbrot (Mandelbrot 1967), where the classic example of fractal dimension deals with measuring the length of the coastline of England that increases when increasing the resolution of the measuring tool and vice-versa. Although natural systems are not truly self-similar nor fractal, the concept of fractal dimension (FD) is useful in a broad range of applications from computer graphics to economy and ecology (Barnsley 2014; Sugihara and May 1990). The main goal of fractal geometry is to describe the variety of natural structures that are irregular, rough, and fragmented (Mandelbrot et al. 1984). Fractal dimension is a unitless number that represents the self-similarity and the complication of a set. Unlike Euclidean dimensions which are expressed with an integer value, the fractal dimension is a fraction that describes the irregularity of objects and how much space they capture. Measuring an object at several resolutions or scales enables estimating the degree of complication and the amount of space filling, as complex shapes tend to reveal more details at higher resolutions. Space filling refers to the ability of objects to reach a higher dimension by filling a region. For example, a line can fill a surface if it passes through every point in it and a surface can fill a volume by folding (i.e., surface complexity). In the case of coral reef 3D structure, both space filling and FD refer to the spectrum between a flat reef and a reef with many structural components of different sizes.

FD can be examined in size categories by calculating the slope between two measurements, (Walsh and Watterson 1993). This provides information not only on niche availability for various taxa but also on the size-frequency distribution of structural features. Therefore, FD is a powerful descriptor of structural complexity (Halley et al. 2004) and important for the scope of coral reef ecology where many of the interactions are size-dependent (e.g., fish use corals for shelter) (Rogers et al. 2014).

Photogrammetry-3D imaging, has emerged as an effective and popular method for benthic surveys (Ferrari et al. 2022). Photogrammetry has been used to measure the rugosity (Friedman et al. 2012), habitat structural complexity (Ferrari et al. 2016), fractal dimension (FD), vector dispersion (Young et al. 2017; Torres-Pulliza et al. 2020; Aston et al. 2022), shelter capacity (Urbina-Barreto et al. 2021; Hylkema et al. 2020) and even to study the impact of storms using structural metrics (Pascoe et al. 2021). The available tools in this field use small (single coral colony) or 2.5D inputs which reduce the z-axis to a single value. Our research deals with reef outcrops (tall and round) that require a new way of analysis: in full 3D on the reef scale. That led us to develop two new methods for calculating the surface complexity of coral reefs using FD. We also calculated four other metrics: surface area over volume (SA/Vol), shelter space (a normalized version of (Hylkema et al. 2020)), vector dispersion (1/K) following (Young et al. 2017), and sphere-packing (Reichert et al. 2017). All of these are methods for calculating surface complexity. We examined each metric separately and also conducted a multivariate analysis as we observed that no single variable was able to fully represent the structural changes of the reef.

To summarize, we employed and analyzed six 3D geometrical metrics (two of which are new) to uncover the changes in structural complexity of seven reef sites in Eilat, Northern Red Sea, over a three-year (2019-2022) period (Fig. 1). We release the ***Reefs4D*** data set, comprising 21 3D models across three time-points: before, immediately after, and two years after a powerful storm. We also release our code for two new FD methods that operate in full 3D and can help in research and monitoring activities. We present a novel analysis framework and an original case study of the effects of an extreme weather event on coral reefs using 3D geometric methods on the reef scale, revealing a trend of resilience in structural components in a two-year time-frame post-disturbance.

**FIGURE 1.**
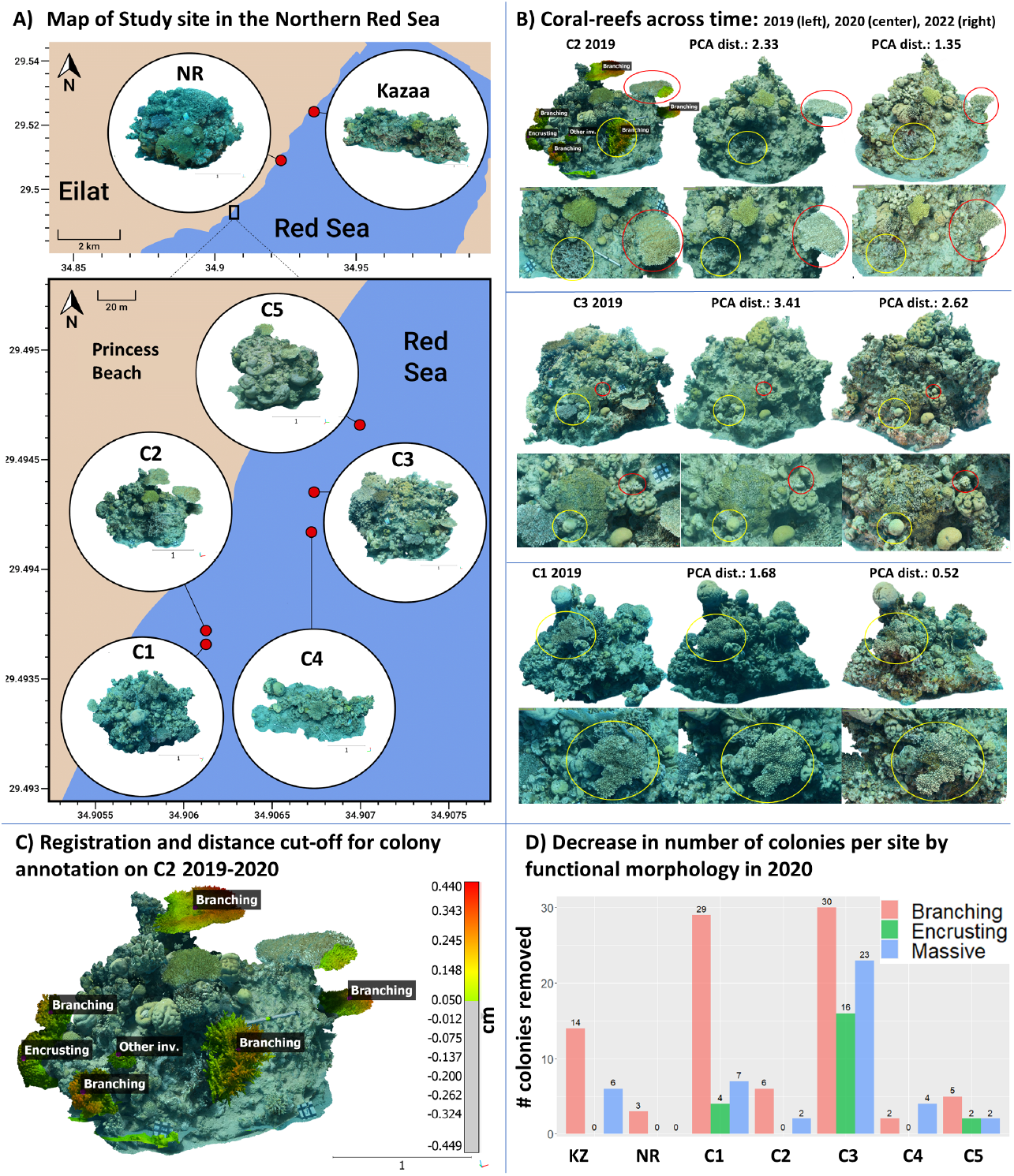
Coral reefs over time. A) A map of the ***Reefs4D*** data set in the northern Gulf of Aqaba (GoA), Eilat. The top map shows the sites Kazaa (KZ, Oil Jetty), Nature Reserve (NR), and Princess Beach (marked by the black rectangle). The bottom map shows sites C1-C5, zooming in on the princess beach area. Each model is shown with a 1 m scale bar. B) Three reefs over time. The PCA distances for each model reflect the changes in structural complexity as shown in Fig. 4C. The circles mark corresponding areas in the models and the close-up views. In C2 (top) the coral colony in yellow is recovering and the one in red is decaying. In C3 (middle), the colony in yellow is static while the one in red is growing rapidly. In C1 (bottom) the colony in yellow is decaying. C) Registration and visualization of C2 in 2019 and 2020. The distance between models was calculated in *Cloudcompare* (CloudCompare 2022) software and a threshold (0.05 m) was used to visualize and count the coral colonies that have been removed from the reef. D) A barplot of the colonies removed by the storm in each reef. The color indicates functional morphology. The majority of corals removed are branching.

## 2 METHODS

### 2.1 3D imaging

Our workflow for 3D imaging is described in (Yuval et al. 2021). We use a DSLR camera and underwater strobes to take overlapping images of the reef from all angles of view and several distances. Primarily, the camera settings and strobes were adjusted to ensure sharp and lit images. Then, using the camera’s intervalometer, subsequent images were captured at one frame per second while swimming around the reef structure and maintaining a stable distance from the substrate. Images were captured with a 50-80% overlap at 24-megapixel resolution in JPG-fine format (Nikon). This was repeated at two swimming distances to the reef (∼ 0.5 m and ∼ 2 m). We used Agisoft Metashape^1^ to process the 3D models by aligning the photos and building mesh from depth-maps. We used high-accuracy image alignment, which was then optimized with “adaptive camera model fitting”. We build mesh from high-quality depth maps with low face count and other processing parameters set to default. The number of images and surface area per model are summarized in Supp. table 1. We scale the models using at least one scale bar placed in the scene. These are objects with distinct features at a known size (e.g., rulers, PVC cards). When scaling the model, we pick the same distinct point in several images, thus reducing the error values below 0.005 m. We then crop the area of interest in the model and export it in PLY format for further analysis. The final resolution of a 3D model is very hard to determine. Based on our previous experiments (unpublished results) and other works in the field (e.g., Figueira et al. 2015) we estimate that our models depict 3D features at the sub-cm scale. We adjust our analysis to take that into consideration by measuring above this size. Since the quality of 3D reconstruction is hard to quantify, we release the *Reefs4D* data set that includes the 3D models from this study to enable further comparisons and evaluation.

**TABLE 1.** Summary of the metrics for structural complexity assessment

| Model | Year | BoxCounting | SpherePacking | 1/K | Alpha/SA | SA/VOI | SA | Vol | ChullVol | Shelter | #Images | Lat | Lon |
| --- | --- | --- | --- | --- | --- | --- | --- | --- | --- | --- | --- | --- | --- |
| C1 | 2019 | 2.190 | 1.669 | 0.799 | 2.295 | 10.848 | 32.663 | 3.011 | 7.151 | 0.579 | 226 | 29.49358 | 34.90619 |
| C1 | 2020 | 2.177 | 1.599 | 0.708 | 2.264 | 10.121 | 22.984 | 2.271 | 4.985 | 0.544 | 265 | 29.49358 | 34.90619 |
| C1 | 2022 | 2.207 | 1.576 | 0.775 | 2.291 | 11.724 | 23.038 | 1.965 | 4.349 | 0.548 | 494 | 29.49358 | 34.90619 |
| C2 | 2019 | 2.198 | 1.541 | 0.763 | 2.262 | 11.990 | 18.921 | 1.578 | 3.491 | 0.548 | 200 | 29.49364 | 34.90618 |
| C2 | 2020 | 2.149 | 1.458 | 0.693 | 2.199 | 10.581 | 12.443 | 1.176 | 2.436 | 0.517 | 226 | 29.49364 | 34.90618 |
| C2 | 2022 | 2.210 | 1.450 | 0.728 | 2.220 | 11.271 | 12.545 | 1.113 | 2.290 | 0.514 | 352 | 29.49364 | 34.90618 |
| C3 | 2019 | 2.263 | 1.726 | 0.876 | 2.283 | 9.152 | 45.358 | 4.956 | 8.710 | 0.431 | 448 | 29.4943 | 34.90678 |
| C3 | 2020 | 2.177 | 1.611 | 0.712 | 2.230 | 7.659 | 22.486 | 2.936 | 5.345 | 0.451 | 159 | 29.4943 | 34.90678 |
| C3 | 2022 | 2.207 | 1.596 | 0.743 | 2.252 | 8.872 | 23.289 | 2.625 | 4.920 | 0.467 | 409 | 29.4943 | 34.90678 |
| C4 | 2019 | 2.154 | 1.488 | 0.805 | 2.265 | 21.799 | 19.499 | 0.894 | 2.755 | 0.675 | 305 | 29.49411 | 34.90677 |
| C4 | 2020 | 2.106 | 1.532 | 0.595 | 2.215 | 11.023 | 18.077 | 1.640 | 3.765 | 0.564 | 196 | 29.49411 | 34.90677 |
| C4 | 2022 | 2.146 | 1.462 | 0.690 | 2.258 | 12.922 | 20.313 | 1.572 | 3.553 | 0.558 | 462 | 29.49411 | 34.90677 |
| C5 | 2019 | 2.205 | 1.511 | 0.823 | 2.307 | 11.470 | 16.540 | 1.442 | 2.678 | 0.462 | 241 | 29.49462 | 34.90703 |
| C5 | 2020 | 2.174 | 1.475 | 0.635 | 2.203 | 8.109 | 11.774 | 1.452 | 2.288 | 0.365 | 204 | 29.49462 | 34.90703 |
| C5 | 2022 | 2.238 | 1.481 | 0.755 | 2.299 | 11.361 | 14.394 | 1.267 | 2.241 | 0.435 | 308 | 29.49462 | 34.90703 |
| Kazaa | 2019 | 2.186 | 1.849 | 0.737 | 2.327 | 10.318 | 89.324 | 8.657 | 21.188 | 0.591 | 832 | 34.93437 | 34.93437 |
| Kazaa | 2020 | 2.181 | 1.840 | 0.747 | 2.293 | 9.599 | 78.811 | 8.210 | 22.276 | 0.631 | 648 | 34.93437 | 34.93437 |
| Kazaa | 2022 | 2.189 | 1.826 | 0.754 | 2.316 | 10.828 | 82.933 | 7.659 | 19.602 | 0.609 | 767 | 34.93437 | 34.93437 |
| NR | 2019 | 2.201 | 1.570 | 0.776 | 2.307 | 9.246 | 32.018 | 3.463 | 7.051 | 0.509 | 392 | 29.50941 | 34.92394 |
| NR | 2020 | 2.184 | 1.633 | 0.802 | 2.203 | 8.327 | 27.164 | 3.262 | 5.610 | 0.419 | 495 | 29.50941 | 34.92394 |
| NR | 2022 | 2.318 | 1.641 | 0.763 | 2.299 | 10.095 | 30.690 | 3.040 | 5.204 | 0.416 | 629 | 29.50941 | 34.92394 |
| Median | NA | 2.189 | 1.576 | 0.754 | 2.265 | 10.581 | 22.984 | 2.271 | 4.920 | 0.517 | 352 | 29.4943 | 34.90678 |
| Std | NA | 0.044 | 0.126 | 0.063 | 0.041 | 2.860 | 23.417 | 2.363 | 6.236 | 0.081 | 195 | 1.9498 | 0.01083 |

### 2.2 Data-set

Our data set contains 21 models of seven shallow reef outcrops at three distinct beaches that represent the different extents of damage following a natural disturbance (Fig. 1A). We refer to them as outcrops because of their pillar shape however, they are fully submerged at depths ∼ 3-8 m. All reef outcrops were smaller than 7 m in lateral extent. The site Kazaa has the largest bounding cube length of 5.95 m, and C5 is the smallest at 2.1 m (after the storm). These represent the typical shallow reef at these locations, consisting of columnar reef structures (outcrops) scattered on a sandy sea bed. The models were captured during Aug-Sept 2019, June 2020, and July 2022. The storm struck Eilat in March 2020. The results are summarized in table 1. We used the models from 2019 as a baseline for all temporal comparisons because they represent an undisturbed state. Furthermore, these models were part of our basic research surveys in underwater photogrammetry. The storm struck these same sites the following year and we identified the opportunity to focus our research on the impact of the extreme weather event on coral reef structural complexity.

### 2.2.1 3D Registration

To enable change-detection and community-level data extraction (coral colony counts) from the 3D models, we conducted 3D registration between the models from 2020 and 2019. By overlaying corresponding models (the same reef over time), the differences can be visualized and help to identify areas of change in the reef (e.g., Fig. 1C). Moreover, registration is important for comparing the results of cube-counting over time because it is a rotation variant method.

However, registration of natural scenes in 3D over time is a challenging problem because it requires static anchor points whereas natural environments are inherently dynamic. Therefore, even manual 3D registration is not always well-defined when conducted on the same reef over time. The models from 2019-2020 were registered in CloudCompare (CloudCompare 2022) using manual registration of at least four anchor points followed by running Iterative Closest Point (ICP) in the software for refining the registration. Anchor points were selected in easily recognizable locations that appear in both models. The results from this registration were used to ground truth an automatic registration method (Alon et al., unpublished results). The automated method was used to register the models from 2019-2022, followed by a manual refinement and inspection in CloudCompare. 3D registration still requires rigorous quality assessment to be used automatically for measuring coral growth and decay. The main challenge is that the direction of change is not constant, i.e., some parts of the reef grow while others decay. To find and annotate the colonies that were removed from the reef, we used a distance cut-off of 0.05 m. The registration and distance color visualization is an effective method for quantifying change detection. Nevertheless, since the models are not fully overlapping, we still inspected both models together to find any other differences that were not highlighted by the distance cut-off.

### 2.3 Structural complexity assessment

We used six geometrical metrics for structural complexity on all 3D reef models. Four known ones: surface area over volume (SA/Vol), shelter-space, vector dispersion (1/K), and sphere-packing, and two new ones that we developed: cube-counting and alpha-shapes/SA.

Fractal geometry has been employed to study coral reefs *in situ* (Knudby and LeDrew 2007), and more recently using Digital Elevation Models (DEMs, where each pixel is a height value) (McCarthy et al. 2022), surface smoothing (Young et al. 2017), and volume-packing (Fukunaga and Burns 2020; Reichert et al. 2017). However, fractal dimension remains challenging to measure in many ecological systems. Measuring it from 2.5D representations (Young et al. 2017) fails to account for overhangs which are a key feature of reefs and provide habitat for many taxa. Box-counting is one of the most common ways of calculating FD (Schroeder 1991). Although there are many implementations of box-counting in 2D, and even on voxel clouds (Fukunaga and Burns 2020) and single coral colonies (George et al. 2021), to the best of our knowledge cube-counting has never been implemented before on wide-scale 3D models of coral reefs.

Past studies either operate in 2.5D or on small 3D inputs (single colony scale). We aimed to push the envelope and conduct a temporal comparison in full 3D on the reef scale using state-of-the-art metrics. Thus, we developed two methods for calculating Fractal Dimension (FD) in 3D: a method for cube-counting on a 3D mesh, and a surface-based method with alpha-shapes (a surface reconstruction method) (Edelsbrunner et al. 1983) and surface area (SA) ratios (alpha-shapes/SA). Measuring SA at different levels of surface details enables us to calculate the dimension, as a flat surface does not decrease in surface area when represented in less detail. The values for fractal dimension measurements range between 2 and 3, where 2 represents a flat surface and 3 represents a volume. We use three methods for calculating FD: cube-counting, sphere-packing, and alpha-shapes/SA because they each portray a slightly different aspect of the overall recovery process. For example, a surface-based method (alpha-shapes/SA) can imply potential resources for grazers, while cube-counting can suggest overall refuge abundance. Sphere-packing is similar to box counting in theory, but the method used here (the software released by (Reichert et al. 2017) is not tailored for reef-scale investigations (it is designed for single coral colonies). Cube-counting is an extrinsic approach that operates on an external 3D grid, therefore, it requires registration to ensure consistency in temporal comparisons. In contrast, alpha-shapes/SA is intrinsic and operates on the mesh surface. Sphere influence is an intrinsic variant of the Minkowski-Bouligand dimension.

### 2.3.1 Cube-counting for fractal dimension

One way to determine the fractal dimension of an object is using the cube-counting method (Schroeder 1991) which is calculated as:

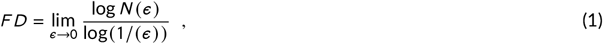

where *ϵ* is the size of the measuring element and N(*ϵ*) is the number of measuring elements required to cover the object. In 3D, the element is a cube, *ϵ* is the length of the cube, and N(*ϵ*) is the number of cubes required to cover the 3D object. This method calculates N(*ϵ*) for several lengths of *ϵ* and evaluates the fractal dimension as the slope of the fitted line between log(N(*ϵ*)) and log(1/*ϵ*).

Our implementation takes an input 3D mesh and bounds it with a minimal bounding cube. The cube is then divided into 8 equal cubes (each dimension is halved). In each iteration, cubes that contain a part of the mesh are counted (1), and only these cubes are then divided in the next iteration (Fig. 2A). In complex shapes the number of cubes in small size-categories is expected to increase exponentially, therefore a logarithmic fit is used. Cube-counting helps to quantify the structural complexity of the reef where degraded reefs are expected to have lower values close to 2, and reefs that are rich in structural elements of different size categories are expected to have values closer to 3.

**FIGURE 2.**
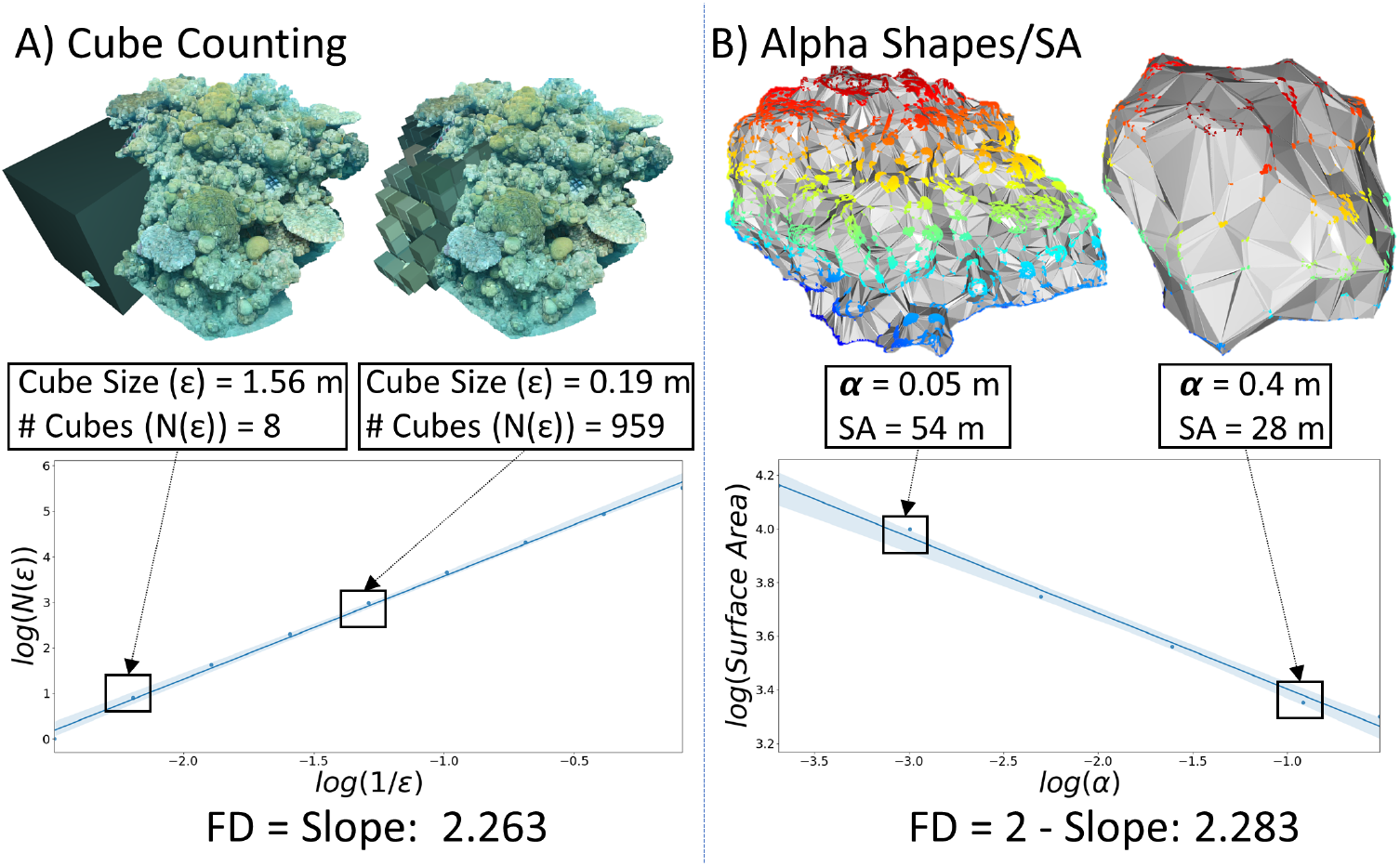
Methods for calculating fractal dimension of coral reefs, demonstrated on model C3. A) The cube-counting method. The displayed box lengths are ½ and 1/16 of the minimal bounding cube which are the 2nd and 5th iterations in cube-counting. Only the corner area of the model is used for the visualization of cube counting. B)The alpha-shapes/SA method shows the alpha-shapes at 0.05 and 0.4 m. Surface area decreases with an increase in alpha.

### 2.3.2 Cube-counting by step-size

The previous analysis can be further expanded to examine size categories. Since reefs are composed of different structural features at multiple scales we wanted to further explore how the storm affected the reef in a size-dependent manner. Measuring FD at several scales is a technique that was mostly used to plot the derivative of the curve (e.g., Fig. 2, blue curves) looking for abrupt changes in slope that indicate a change in structural regime (Walsh and Watterson 1993; Backes and Bruno 2012; Nash et al. 2013). We analyzed the FD per size category using the cube-counting method. Determining the FD per cube size is done by measuring the slope between two points in the cube-counting results (Eq. 1, e.g., Fig. 2A).

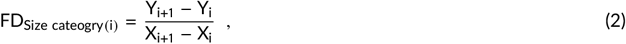

Where *Y*_*i*_ = log(*N* (*ϵ*)) is the number of cubes necessary to cover the reef at size category *i* where the element size is *ϵ*, and *X*_*i*_ = log(1/*ϵ*).

To calculate change per size category, we compare each measurement to the result of the 2019 measurement by subtraction so that negative values indicate a decrease in FD and vice versa. There are slight differences in box sizes when comparing models of the same reef over time because they depend on the size of the original bounding cube (Supp. Fig. 7). Therefore, the size values in the step size analysis are the average size per category. The size of the original bounding cube depends on the model’s size and orientation, making the registration is important for this calculation. Larger variation occurs in the large-size categories, whereas the values converge in the small-size categories. Therefore, we only examine the last five iterations of cube-counting. For example, in the size category of 40 cm, we are dealing with the third iteration of cube-counting (in models C1, C3, C4, NR) where each box has more influence on FD, causing increased variation and sensitivity to rotation and registration.

### 2.3.3 Alpha-shapes and surface area

Alpha-shapes (Edelsbrunner et al. 1983) is a surface reconstruction method that generalizes the concept of a convex hull of a set of points. In simple terms, the alpha-shape algorithm takes a set of points in space and constructs a shape that encapsulates them. This shape is obtained by defining the parameter alpha (*α*), which determines the level of detail in the shape. A small alpha value will result in a shape that closely fits the points, while a larger alpha value will result in a more simplified shape. From an ecological perspective, this method smooths out features that are smaller than alpha, thus enabling the examination of the reef at different resolutions.

In the method we developed we look at how the surface area of the alpha-shape changes with *α*. In general, we expect the surface area to decrease when *α* increases, as then the shape is smoother. Flat and smooth shapes will not decrease in surface area when increasing alpha as opposed to complex and rugged shapes, since the alpha filter smooths out their structural features. Our insight is that the rate of this change (surface area to alpha) can indicate the FD.

For *α* = (0.025, 0.05, 0.10, 0.2, 0.4, 0.6) meters, we calculate (example in Fig. 2B):

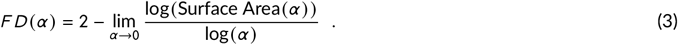

We chose to start from 0.025 m to ensure sampling above our model resolution, and that the results are not distorted by the alpha filter which on small numbers creates cavities in the mesh and increases the SA metric disproportionately. We stop at 0.6 m because at this alpha value we almost reach the convex hull of the full set of points which is the most crude shape with the lowest SA. The results range from 2 to 3 where higher values indicate more surface ruggedness. To calculate the alpha-shape and surface area, we used the Open3D python package (Zhou et al. 2018).

### 2.3.4 Sphere influence for fractal dimension

This measure calculates the fractal dimension using a variant of the Minkowski–Bouligand method- the sphere packing method. It places spheres of increasing radius on the mesh vertices and measures the volume of the shape at each iteration as the influence volume. The ratio between sphere size and its influence (i.e., the overall volume) defines the fractal dimension. To calculate this measure we used the software and guidelines provided by the authors in (Reichert et al. 2017). The main disadvantage of their software is that it is designed for single coral colonies and it stops at an upper limit of 20 cm of sphere influence.

### 2.3.5 Surface Area (SA) / Volume

In this study, we refer to SA/Volume as a single measure. Surface area and volume were calculated in Metashape using the built-in *view mesh statistics* tool. The surface area was calculated first. Then all holes in the mesh were closed to 100% and volume was measured. Dividing surface area by volume provides a notion of the amount of surface complexity, with higher values for more complex structures and lower numbers for flatter surfaces.

### 2.3.6 Shelter-space

The shelter-space metric quantifies the amount of volume available for motile reef organisms in the immediate vicinity of the reef. We calculate the convex hull of the mesh in the open-source software Meshlab (Cignoni et al. 2008). We divide the volume of the 3D mesh by the volume of the convex hull to obtain the shelter-space metric using the following formula:

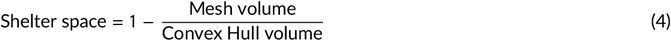

This metric ranges from 0 to 1 where 0 is no shelter space and higher values mean more shelter space.

### 2.3.7 Vector dispersion 1/K

This metric is an estimate of reef complexity derived by measuring the angles of the normals of the mesh faces. The result ranges between 0 to 1 indicating increasing complexity with increasing values, with flat surfaces closer to zero. We calculated the direction cosines (cosine of the angle between the normal and each one of the axes) of each face in the mesh using the Open3D package (Zhou et al. 2018) and a custom Python script following (Young et al. 2017):

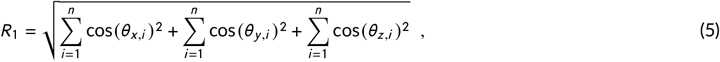

where *θ*_*x,i*_, *θ*_*y,i*_, *θ*_*z,i*_, are the angles between the ith normal and the x,y,z axis, accordingly, and n is the number of faces in the mesh

Then, the Vector Dispersion measure 1/*K* is:

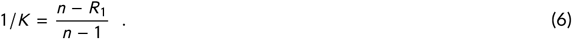

Vector dispersion is useful for estimating the surface complexity of the reef because flat surfaces have an *R*_1_ value close to n (the number of faces) resulting in a very small fraction close to zero (Eq. 6).

### 2.4 Statistical Evaluation

We studied seven reefs at three time points using six metrics. To evaluate the demise and recovery of the reef, we compared the result of each measurement to the baseline from 2019. We used a non-parametric statistical test because our data set was small and did not meet the assumptions of parametric tests. We used a one-sided paired samples Wilcoxon test (n_pairs_ = 7) to compare the difference in each variable from the baseline in 2019 (Fig. 3B). We tested whether the differences between 2019 and 2020 were more negative (smaller values) than the differences between 2019 and 2022. The difference was significant for cube-counting, alpha-shapes/SA, and SA/Vol with P-values of 0.0078, 0.0111, and 0.0078. The results were not significant for sphere-packing, 1/K, and shelter space (P-values were 0.9609, 0.05469, and 0.4687).

**FIGURE 3.**
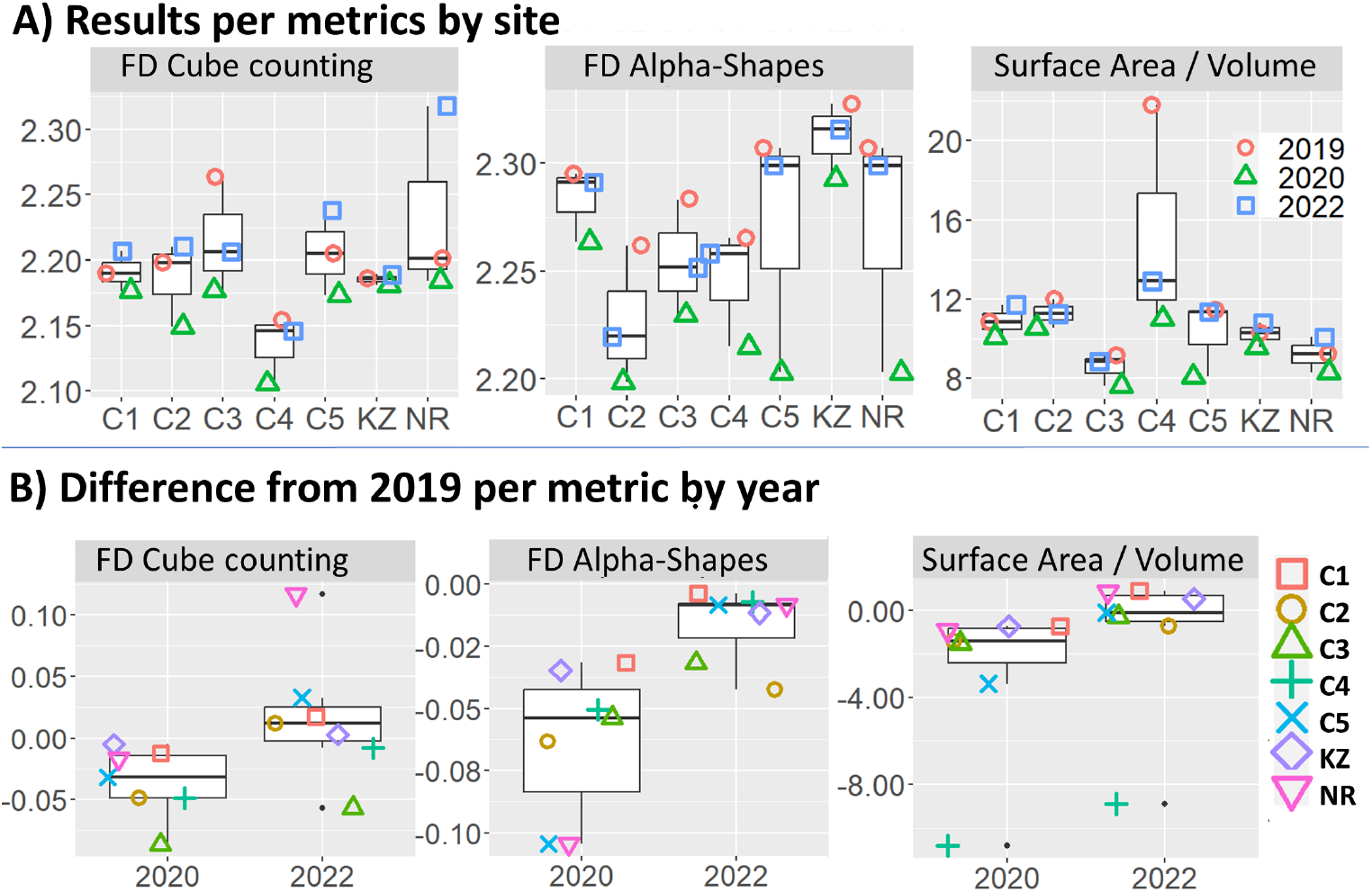
Metrics for structural complexity assessment. A) Results of cube-counting, alpha-shapes/SA, and SA/Vol on all models. Color and shape indicate time. The medians are summarized in Supp. Table 1. B) The difference in metric per year with the 2019 models as a reference. Color and shape indicate the site. We used a paired samples Wilcoxon test (n_pairs_ = 7) with a one-sided alternative to test if the difference in each metric in 2020-2019 is smaller than the difference in 2020-2019. The difference was significant for cube-counting, alpha-shapes/SA, and SA/Vol with P-values of 0.0078, 0.0111, and 0.0078. The rest of the metrics are shown in Supplementary Figure 1.

Testing each metric separately is very difficult because some metrics exhibit a recovery or even increasing values, while for the same site and year, a different metric may show a decrease (e.g., cube-counting and shelter space on C3 2020, Fig. 3A). Combining the different variables helps to see how each site was affected and to reveal a complete picture of changes in the reef. To explore the relations between variables we calculated Pearson’s correlation coefficient by looking for highly correlated pairs (Fig. 4A). We performed a Principal Component Analysis (PCA) to reduce the dimensionality of the data (Fig. 4B). The third principal component (PC3) explains less than 10% of the variance in the data and was excluded from the visualization. This enabled us to represent each sample (model per time) as a single data point in two dimensions. For each point from 2020 and 2022, the Euclidean distance to the corresponding baseline point (2019 from the same site) was measured. This provided an overall view of the changes in the reef considering all six metrics.

**FIGURE 4.**
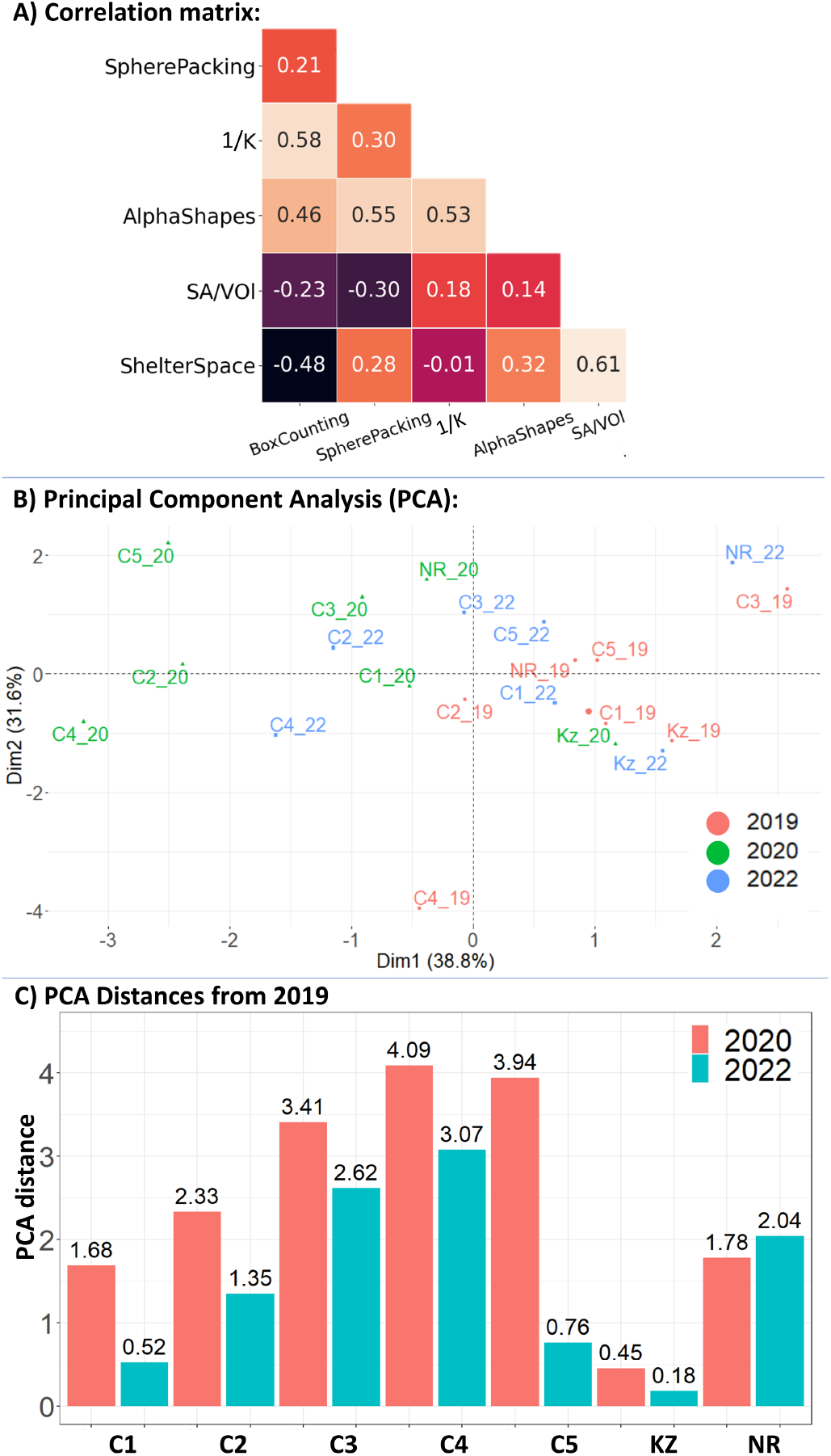
Multivariate analysis of coral reef structure over time. A) Pearson product-moment correlation coefficients of all metrics. The figure shows a low-to-moderate correlation among the variables. B) Principal component analysis (PCA) of the full data set. Colors signify the year and reveal a clear temporal trend along PC1. C) Euclidean distance on two PC axes between years per site, with the 2019 models as reference. The color indicates the year. The distances are shorter in 2019-2022 (blue) than in 2019-2020 (red). The red bars correspond with measurements taken right after the disturbance and depict a large difference from the baseline. The blue bars are shorter than the red bars in almost all of the sites, indicating that there is a recovery in structural complexity.

### 2.5 Code and data availability

Code will be made available at: https://github.com/MatanYuval/ReefMetrics

Data will be made available at: https://zenodo.com/

The reader is encouraged to view the models in 3D online at https://skfb.ly/oGJDB.

## 3 RESULTS

Overall, we found that there is more difference between the years 2019 and 2020 than between 2019 and 2022, i.e., two years after the extreme weather event, suggesting a recovery in structural complexity. This was significant in several individual metrics and further exemplified by our multivariate and step-size analysis.

### 3.1 Visual change

The storm caused such pronounced damage to the reef that when comparing the models from 2020 to 2019 we were able to count the number of coral colonies removed by the storm using manual registration and a mesh-to-mesh distance visualization (CloudCompare 2022 example in Fig. 1C). We found that the majority of corals removed (broken) by the storm were those with branching morphologies (Fig. 1D) which was probably due to their life-history traits and fast-growing fragile forms.

Comparing models from 2022 with those from 2019 to quantify coral growth was much more challenging. Primarily, the registration was more difficult due to added changes over time. Second, the signal (coral growth) is much more mild compared to that of a full colony removal. Moreover, tracking corals over time in 3D is not straightforward. We observed that show that while some corals grow others decay, and even the same coral colony can show reverse trends. The close-up views in Fig. 1B show several examples. In model C2 (Fig. 1B, top) the coral colony in yellow is recovering and the one in red is decaying. In the model C3 (Fig. 1B, middle), the colony in yellow is static while the one in red is growing rapidly. In C1 (Fig. 1B, bottom) the colony in yellow is decaying.

### 3.2 Individual metrics for structural complexity assessment

Three measures (cube-counting, alpha-shapes/SA, and SA/Vol) were indicative individually of decline and recovery in structural complexity (Fig. 3). The other metrics were not significant by themselves, but they still contributed to the multivariate analysis. Cube-counting and alpha-shapes/SA values were in the expected ranges of 2-3. The results of sphere-packing are in the range of those shown by the authors of this method (Reichert et al. 2017) which are slightly lower than 2. In cube-counting in all models the results dropped between 2019 and 2020 and then increased again in 2022. Interestingly, in most reefs, the measure in 2022 is higher than the one in 2019. In alpha-shapes/SA and SA/volume, all numbers dropped in 2020, then increased in 2022, but did not go beyond the 2019 measure. SA/Vol and shelter space have extreme values for C4 in 2019 that are caused by a large footprint of the 3D model which was reduced after the storm. When looking at the other three metrics that were not significant (Supp. Fig, 6A), there is a temporal trend where the 2019 measurement of structural complexity (1/K, sphere packing, and shelter space) is generally the highest and 2020 is the lowest. There are reverse trends between years for some of the metrics, for example for site C3 we have the lowest values in shelter space and 1/K, and the highest sphere packing values. In site NR the trend is opposite, as it has the lowest values in sphere-packing and the highest values for 1/K. When looking at the differences from 2019 in vector dispersion 1/K (Supp. Fig, 6B), there is a trend in recovery for the Princess beach sites (C1-C5) but not for the NR and Kazaa (KZ) sites. These results emphasize the importance of using more than one metric in studies of structural complexity in the benthos.

### 3.3 Multivariate analysis

We calculated the Pearson correlation coefficient between variables (Fig. 4A). All variables had medium-to-low correlation (-0.48 to 0.61). This means that they are not measuring the same characteristic and demonstrates the importance of using more than one metric in describing the 3D structure of coral reefs. We found that shelter space had the highest correlation with SA/Vol, followed by box counting and 1/K, sphere-packing and alpha-shapes, and 1/K and alpha-shapes. 1/K and shelter space had the lowest correlation. Shelter space and SA/Vol both include the SA component in their calculation which can help to understand why they are highly correlated. 1/K and alpha-shapes are both intrinsic methods that operate directly on the mesh which may contribute to their correlation. Box counting and alpha-shapes/SA have a correlation of 0.46, which helps to show that different FD methods contain different ecological information.

When examining the PCA results (Fig. 4B), we found that PC1 correlates with 1/K, sphere-packing, and alpha-shapes. These are all intrinsic methods that operate on the mesh directly. Practically, intrinsic methods operate without an external coordinate system and are not sensitive to rotation making them easier and more robust to employ in temporal comparisons. In contrast, PC2 correlates with SA/Vol and shelter space, both of which include the mesh volume in their calculation. The ordination data points indicate a temporal trend of recovery mostly along PC1. Notably, the site Kazaa (KZ in Fig. 4B) remained clustered which matches our visual observation that it was affected the least and degraded the least.

3.3.1 PCA distances

We measured the absolute distances of points on the first two PCA axes within the site and across time, comparing them to the 2019 models as a baseline. The results of these measurements are presented in Fig. 4C. This analysis emphasizes the overall change per site and provides information on how certain sites were clustered within the PCA analysis. The site KZ (Kazaa) shows the least change from the baseline, both immediately after the storm (red) and also in recovery (blue). This was also observed visually. Sites C3 and C4 and C5 suffered the most damage (Princess Beach sites). C3 and C4 are still far from their baseline state although they are starting to recover (the blue bar is shorter than the red bar). The sites C1 and C5 (Princess Beach) were damaged by the storm, but their recovery in structural complexity is substantial as they are very close to the baseline (the blue bar is very short). The shorter distances in 2019-2022 (red, Fig. 4C) generally indicate a positive trend of recovery and increasing structural complexity. The extracted distances (Fig. 4C) can be examined together with the PCA to provide more insight. For example, for the site NR, the blue bar is longer than the red, meaning that it is still different from the baseline. When looking at the PCA results Fig. 4B) it seems that this site has increased in structural complexity in comparison to the baseline. We can infer this because we found that this PC axis (PC1) indicates an increase in structural complexity and NR22 is further to the right than NR19.

### 3.4 Comparison of FD by size-category using cube-counting

In Fig. 5A we depict the FD per size category, shown as the difference from the 2019 measurement. We found that there was a decrease in FD following the storm in all sites in all size categories except for C4 and C5 (5A, blue bars), where there is an increase in FD on the 5th size category (∼ 40 cm). This increase can be an artifact of registration as this size category is the second to the third iteration of cube-counting in most models (large models like KZ start from a larger bounding cube) and a slight change in the model’s orientation can lead to a different number of boxes needed to cover it. The differences between 2022 and 2019 (5A, orange bars) show a trend of returning to the baseline almost in all size categories and all sites, although in some cases we see a further decrease, such as in C1, C2, and KZ on the 20 cm category.

**FIGURE 5.**
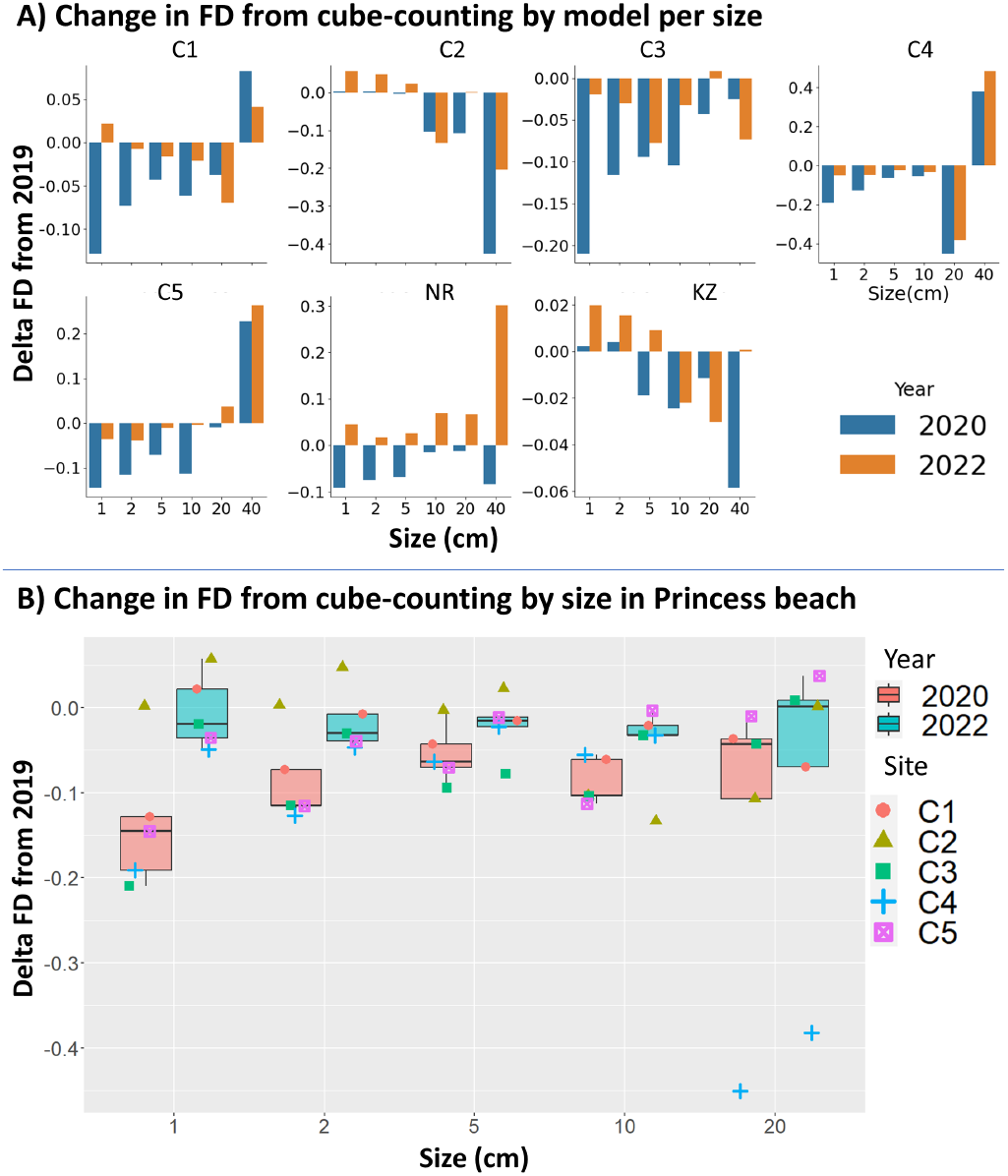
Comparison of FD by size-category using cube-counting. A) The changes by size category (cube size) per site compared to 2019 as a baseline. We calculated the slope between two consecutive measurements in cube-counting and attributed the value to the smaller box size (see methods 2.3.2). B) The changes in Princess Beach sites (C1-C5) by size category. The sites are marked by shape and color, and the years are aggregated in the boxes. The blue boxes are higher than the red boxes showing that there was more difference (decline) in FD in 2020 than in 2022.

**FIGURE 6.**
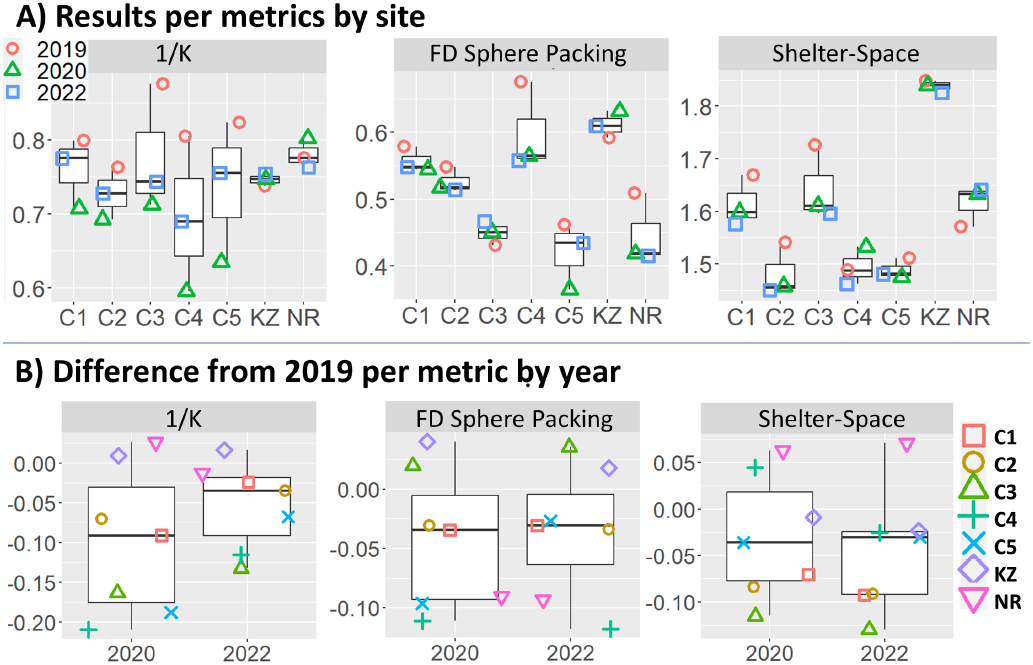
**Supplementary figure 1:** Metrics for structural complexity assessment. A) Color and shape indicate time. The medians are summarized in Table 1. B) The difference in metric per year with the 2019 models as a reference.

**FIGURE 7.**
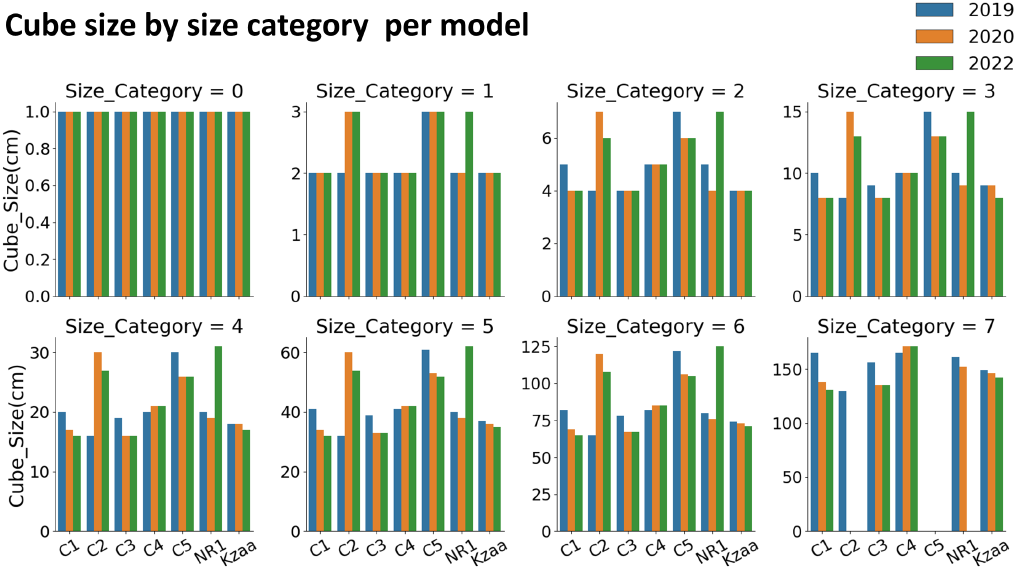
**Supplementary figure 2:** Size of cube length from cube counting by step size in cm per all models by size categories. Color signifies the year. The smaller size categories converge and the large size categories have more variation. the standard deviations of the different size categories are: 0, 0.46,1.12, 2.52, 5.04, 10.02, 19.85, 14.46

We then aggregated the size categories and plotted the change per step size at Princess Beach (sites C1, C2, C3, C4, and C5) using 2019 as a reference (Fig. 5B). We chose to focus on these sites because they are located on the same beach and were the most exposed to the storm. The results show that the smaller size categories exhibit more recovery which can be attributed to coral growth. Overall, there is less difference between the years 2019 and 2022 (red boxes) than between 2019 and 2020 (blue boxes), i.e., two years after the extreme weather event, suggesting reef resilience: the reef is returning to its former cube-counting dimension.

## 4 DISCUSSION

Complex dynamic systems are governed by different processes that occur simultaneously on different levels of selection (Okasha 2006). Understanding them separately as well as their interplay is one of the key goals in ecological research (Green et al. 2006).

In coral reefs, structural complexity emerges from lower-level interactions between reef organisms (Jackson and Hughes 1985). To a large part, coral life histories (e.g., growth rates and morphologies) and community-structure (e.g., size-frequency distributions, species composition) dictate the shape of the reef (Jones et al. 1994). The 3D structure is formed over large periods, retains memory, and encapsulates ecological information. Thus, it can be useful as a descriptor of ecosystem alternate states, and an indicator for shifts between them. At the same time, the concept of coral reef structural complexity has many different components relating to different ecological processes. Therefore, it is important to use various structural metrics to gain a holistic view of the ecosystem state and trajectory. Moreover, underwater image collection is less replicable than a standard lab protocol. It is difficult to capture the exact same images from the same angles over time and external factors such as light (e.g., cloud cover) and sea conditions heavily influence the image quality, especially in shallow waters. Using several structural metrics helps to buffer out noise or bias that originate from the 3D reconstruction process.

Here we used a combination of 3D geometrical metrics to paint a detailed picture of the effect of an extreme weather event on the structural dynamics of coral reefs. The storm in 2020 is the most severe to hit the region in several decades. The reefs in 2019 had ample time to develop over the decades preceding the storm. They represent an evolved state with regard to structural complexity and were used as a pre-disturbance baseline. The survey in 2020 was conducted several weeks after the storm, depicting the reef immediately (in terms of the coral community) post-disturbance. The 2022 surveys were far enough apart from the storm to enable capturing coral growth and recovery. We aim to continue collecting 3D models of these reefs in the upcoming years for further research on coral community dynamics and structural complexity. Comparing the results to the baseline shows that the reefs are returning to their pre-disturbed state. This is especially apparent in the multivariate analysis. Coral reefs operate in an ecological latent space where peaks are tipping points and valleys are stable states. Resilience relates to the transitions between them. Although structural complexity represents only a fraction of this space, we were able to use it as a practical descriptor. It enabled the depiction of a disturbed state and a partially recovered state using structural metrics, which is extremely important considering the role of structural complexity in maintaining coral reef functions. Our results show that the reefs have not completely diminished due to the storm. In contrast, they are recovering in structural complexity, i.e., the effect of the storm was reversible. Although resilience was not directly measured, we managed to gain an interesting angle on it in terms of structure, i.e., structural resilience. Coral reef resilience includes many more functional levels which were not covered in this study. Nevertheless, we add a piece to the puzzle towards building a new picture of reef resilience and the contribution of different components of habitat structural complexity.

Not all reefs were equally impacted by the storm nor reacted in the same way. The southernmost sites (Princess Beach, C1-C5), were the shallowest (∼ 3 - 6 m) and closest to the slope of the reef, thus exposed to higher wave energy and affected the most. The reef at Kazaa was the largest in our study, being elongated, very complex, and relatively sheltered by its location. This site seemed to change the least. The reef at NR was the deepest (∼ 8m) and was not severely affected.

Fractal dimension in reef ecology has been measured in various ways for several decades (Bradbury et al. 1983). However, these measurements were largely confined by the available technological tools. The new era of underwater photogrammetry (Ferrari et al. 2022) unleashes the opportunity to study reefs in 3D, in accuracy and detail across large scales, driving FD to become one of the most common descriptors for coral reef ecology. We used three methods to calculate FD: cube-counting, alpha-shapes/SA, and sphere-packing (Reichert et al. 2017). As opposed to previous works, our implementations for calculating FD (cube-counting and alpha-shapes/SA) leverage the full 3D model rather than a 2.5D representation (single Z value per coordinate) and perform on the entire reef scale rather than single coral colonies. This is important in light of the input: reef outcrops, which are tall, round, and contain overhangs and crevices. Many protocols for underwater photogrammetry are based on downward-looking imaging for large-scale surveys (e.g., 100 m2 transects) and thus do not lose ecological information from this reduction. However, depending on the input and especially in small, single reef reconstructions (an order of magnitude smaller than a standard monitoring plot), it is advantageous to generate a comprehensive 3D reconstruction and analyze it appropriately-in full 3D. Cube-counting and alpha-shapes/SA are effective methods that were indicative individually of decline and recovery in structural complexity (Fig. 3). The main disadvantage of cube-counting is that it is an extrinsic calculation. This means it is sensitive to rotation and registration errors since the minimal bounding cube (the first iteration of the algorithm) is axis-aligned on an external grid. An interesting follow-up study would focus on finding a rotation-invariant minimal bounding cube. That would eliminate the need for registration in temporal comparisons. The sphere-packing method is promising as an intrinsic calculation of FD and an interesting follow-up study would develop an implementation for wide-scale 3D models. We chose to introduce a new surface-based measure using alpha-shapes because it enables calculating FD in 3D and uses a size-based filter that can be interpreted meaningfully. Alpha-shapes/SA is an intrinsic method that is not sensitive to rotation because it operates on the model directly without considering the external coordinate system (grid). A disadvantage is that sometimes the alpha filter causes distant areasof the mesh to merge together resulting in a reverse ratio (low alpha value with low surface area).

Calculating FD by size category from cube-counting enables choosing the scales of investigation (Figs. 2, 5) (Backes and Bruno 2012). Defining the relevant contribution of each size class to an observation, i.e., structural complexity measurements, can lead to a better understanding of the overarching process. For example in our case, we attribute the decrease in FD in large size-categories to whole colony removal by the storm, and an increase in the small size-categories is attributed to coral recruitment and growth. For example, in C2, C3, C4, and KZ, the decline in FD was at a size above 5-10 cm whereas in sites C1 and C5 the decrease was on small size categories. An interesting follow-up study would focus on comparing the size of corals that were dislodged by the storm with the structural complexity of the substrate and the size of the reef outcrop. Shelter space and SA/Vol are sensitive to the models’ footprint (the size of the model’s base, the area that the reef captures on the sand/substrate), especially shelter space which uses a convex hull function (crude encapsulation of the model). For example, these metrics have extreme values for C4 2019 that are caused by a large footprint of the 3D model which was reduced after the storm. Defining the outline of the model is not straightforward as the surrounding environment is also changing, especially in extreme storms. We chose to crop the reef immediately above the substrate in order to facilitate the registration process. Model registration is important for visualizing changes in the reef as well as for calculating rotation variant methods (extrinsic methods, e.g., cube-counting). However, registration is difficult because the scenes are dynamic. The reef and the substrate can be tilted by the storm, and both of them can even become tilted in opposite directions. External coordinates (e.g., differential GPS) could help to register the models over time yet they are difficult to obtain underwater. An important follow-up study would focus on defining the error of registration in practical terms, i.e., the amount of growth/decay (signal) which can be captured. This can be done by imaging the same reef several times on the same day and registering the models assuming there is no change (Figueira et al. 2015; Yuval et al. 2021).

One of the main shortcomings of this study is the variance in the number of images collected in each survey. Ideally, we would have a consistent amount of images per surface area depicted. However, we wanted to guarantee a high-quality reconstruction of the reef. Therefore, the third survey round had a larger number of images per site and a higher-quality reconstruction which affects the complexity score (more complex). We solve this in cube-counting and alpha-shapes by measuring above the size of reconstruction artifacts: the smallest bounding cube is one cm^3^ and the smallest alpha value is 2.5 cm.

Coral reefs are some of the most intricate and diverse ecosystems on the planet, and the problem of scale vs. detail in coral reef ecological studies has yet to be brought to bear. The most important follow-up study will measure coral growth and decay directly by tracking the same colonies over time (Fig.1B). Although there are tools for fast segmentation of benthic maps (Pavoni et al. 2022), there are almost no available tools for 3D segmentation and accurate size measurements in complex 3D scenes (Petrovic et al. 2014). 3D segmentation tools still require manual annotations that limit their scalability. On the contrary, using only geometry is scalable but provides limited detail, i.e., lacking taxonomic information. The models in our data set contain thousands of single coral colonies and counting them separately on each model is rigorous and not optimal. We opted for 3D registration to help us focus our attention on areas of the reef where colonies were removed. However, a thorough analysis of each model separately will enable much richer data extraction.

In light of recent advances in deep learning, (e.g., Kirillov et al. 2023), we expect that in the near future, 3D instance segmentation (a deep-learning task that segments every object in a model) and geometric deep-learning will be employed in studies focusing on image-based ecology. Moreover, with advances in computation, we expect that the role of simulations will become more dominant in ecological studies. We release the ***Reefs4D*** data set, which contains detailed 3D models of real complex systems that are scarce in the computer-vision and 3D graphics community and can help to solve the aforementioned challenges and lead to exciting research.

Corals dominate many shallow marine habitats and powerful storms are inseparable from their evolution. Nevertheless, such acute disturbances can cause coral reefs to become more vulnerable to local disturbances (e.g., pollution, tourism), which in turn can lead to a catastrophic ecosystem collapse. In light of the increasing frequency of disturbances, it is important to continue tracking reefs over large areas and long periods.

## Declarations

### Author Contributions

**All authors:** Conceptualization, Writing—Review and Editing. **M.Y**., **N.P:** Software, Methodology, Visualization. **M.Y**.: Formal analysis, Investigation, Data Curation, Writing—Original Draft. **T.T:** Supervision.

#### Acknowledgments

We thank the Interuniversity Institute for Marine Sciences of Eilat for making their facilities available to us. We thank Aviad Avni and Opher Bar-Nathan for valuable intellectual and technical contributions; Netta Kasher for drawing Fig. 1A; We thank the reviewer for the constructive comments. Fieldwork in Eilat was carried out under permit 42-128 from the Israeli Nature and Parks Authority.

## Supporting Information

The results of structural complexity calculations, the number of images, and the location of the models.

## Footnotes

1 Agisoft Metashape Professional (Version 1.5-1.7) (Software) Agisoft LLC, St. Petersburg, Russia, 2016.

